# Measuring the neurodevelopmental trajectory of excitatory-inhibitory balance via visual gamma oscillations

**DOI:** 10.1101/2024.09.10.612351

**Authors:** Natalie Rhodes, Lukas Rier, Krish D. Singh, Julie Sato, Marlee M. Vandewouw, Niall Holmes, Elena Boto, Ryan M. Hill, Molly Rea, Margot J. Taylor, Matthew J. Brookes

## Abstract

Disruption of the balance between excitatory and inhibitory neurotransmission (E-I balance) underlies theories of many neurodevelopmental disorders, however, its study is typically restricted to adults, animal models and the lab-bench. Neurophysiological oscillations in the gamma frequency band relate closely to E-I balance, and a new technology – OPM-MEG – offers the possibility to measure such signals across the lifespan. We used OPM-MEG to measure gamma oscillations induced by visual stimulation in >100 participants, aged 2-34 years. We demonstrate a significantly changing spectrum with age, with low amplitude broadband gamma oscillations in children and high amplitude band limited oscillations dominating in adults. We used a canonical cortical microcircuit to model these gamma signals, revealing significant age-related shifts in E-I balance in superficial pyramidal neurons in visual cortex. Our findings detail the first MEG metrics of gamma oscillations and their underlying generators from toddlerhood, providing a benchmark against which future studies can contextualise.

## Introduction

The maintenance of a balance between excitatory and inhibitory neurotransmission (E-I balance) is essential for healthy brain function and its disruption underlies a range of psychiatric conditions, notably autistic spectrum disorder (ASD) (Nelson and Valakh, 2015; Rubenstein and Merzenich, 2003; Sohal and Rubenstein, 2019). High frequency neurophysiological oscillations in the gamma range (>30 Hz) play a key role in information processing (Fernandez-Ruiz et al., 2023) and arise due to interactions between neuronal excitation and inhibition (Bartos et al., 2007; Vinck et al., 2013). Thus, measurement of gamma oscillations can provide a powerful metric of E-I balance (Gray et al., 1989; Gray and Singer, 1989; Whittington et al., 1995). Despite this importance, our understanding of gamma oscillations, their developmental trajectory in early childhood and perturbation by disorders remains poorly characterised and this is largely due to instrumental limitations. Here, we use a new neurophysiological imaging platform to measure gamma oscillations in individuals from early childhood to adulthood and use a model of neural circuitry to investigate their underlying neural generators.

Gamma oscillations can be measured non-invasively using either electro- or magnetoencephalography (EEG or MEG), with MEG providing more robust data. However, both techniques have limitations, particularly for children. In EEG, the gamma signal (which manifests as an electrical potential difference across the scalp surface) is diminished in amplitude and distorted spatially by the skull (Baillet, 2017). EEG gamma signals are also obfuscated by interference generated by non-neural sources such as muscles (Boto et al., 2019; Muthukumaraswamy, 2013) making it difficult to measure gamma reliably, particularly if subjects move (which is common in children). MEG, which measures magnetic fields generated by neural currents, is less affected by non-neural artefacts and has better spatial specificity than EEG (because magnetic fields are less distorted by the skull than electrical potentials). This means that gamma oscillations have a higher signal-to-noise ratio (SNR) and their origin can be better localised when using MEG rather than EEG (Muthukumaraswamy and Singh, 2013). Multiple studies argue that MEG is the measurement of choice for gamma oscillations (Gaetz et al., 2011; Hall et al., 2005; Muthukumaraswamy et al., 2010, 2009; Orekhova et al., 2015; Takesaki et al., 2016; Tan et al., 2016). However, MEG systems classically rely on cryogenically cooled sensors that must be fixed in position in a one-size-fits-all helmet. Such systems cannot cope with changing head size through childhood or large subject motion relative to the (static) sensors. Consequently, most extant MEG studies of gamma oscillations are limited to adults.

As ASD has a typical diagnostic age of 3 years and above, if we are to understand its neural substrates, E-I imbalance (and gamma oscillations) must be measured reliably in children from <3 years of age and upwards. To this end, we measured the developmental trajectory of gamma oscillations using a new technology – optically pumped magnetometers (OPMs) (for a review see Schofield et al., 2022). OPMs uniquely allow MEG signals to be recorded using wearable helmets (Boto et al., 2018,; Hill et al., 2020), which adapt to different head sizes and enable movement during scanning. This provides an ideal environment to gather high fidelity data in children, and studies have already shown that OPM-MEG can be used to measure neurophysiological signals in the early years of life (Corvilain et al., 2023; Hill et al., 2019) and that neurodevelopmental changes in neurophysiology can be assessed (Rier et al., 2024). This platform therefore offers the best chance for measurement of gamma oscillations and subsequent modelling of neural circuits, to understand how E-I balance changes with age.

We characterised the neurodevelopmental trajectory of gamma oscillations from age two years to adulthood. We used a newly developed child-friendly OPM-MEG system to collect data during a visual task that is known to elicit gamma oscillations in primary visual cortex (Orekhova et al., 2018). These visual gamma effects have been associated with feature integration (Eckhorn et al., 1988; Gray et al., 1989), object representation (Tallon-Baudry and Bertrand, 1999), and selective attention (Fell et al., 2003). Existing studies suggest these oscillations are altered in childhood (Gaetz et al., 2011; Orekhova et al., 2018) (albeit in older children), in ASD (Orekhova et al., 2023), and twin studies suggest they are highly heritable, having a strong genetic component (Pelt et al., 2012). The cellular generators of visual gamma oscillations have been described (Spaak et al., 2012; Xing et al., 2012) by modelling the interaction between superficial pyramidal cells and inhibitory interneurons within V1. Similarly, we use a dynamic causal model (DCM) – based on a canonical cellular microcircuit (Shaw et al., 2017) – to investigate the contributions of inhibitory and excitatory neurotransmission to the gamma signal. We hypothesised that OPM measurement of gamma oscillations alongside DCM would demonstrate an E-I balance change as the human brain matures.

## Results

OPM-MEG data were collected using either a 192-channel system (located at the Sir Peter Mansfield Imaging Centre, University of Nottingham, UK (UoN)) or an 80-channel system (located at SickKids Hospital, Toronto, Canada (SK)). The two systems had a similar design (Figure 1a; Cerca Magnetics Ltd. Nottingham, UK) and channels were located to ensure good coverage of the visual cortices. (See also supplementary information (SI) Table S1; Equivalence between systems is shown in Figure S1.)

**Figure 1.**
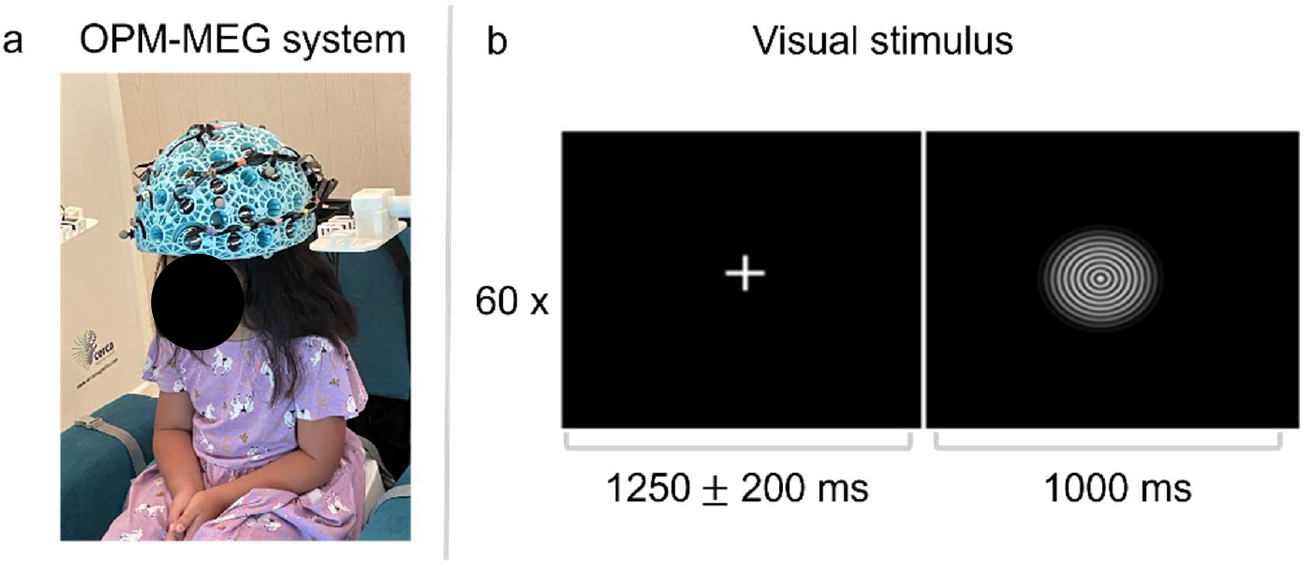
Methods. a) An image of a child in the OPM-MEG system, b) the concentric circles visual stimulus and paradigm timing, which was presented for 60 trials.

102 typically developing participants (aged 2 – 34 years; 44 male; see SI Table S2) took part in the study, which was approved by ethics review boards at both sites. Participants viewed visual stimuli comprising inward drifting circular gratings (moving at 1.2°*s*^−1^) (Figure 1b). A single experimental trial started with a white fixation cross located centrally on a black screen for 1.25 ± 0.2 s. This was followed by 1 s of stimulation. 60 trials were recorded per subject, and trials were interspersed with pictures of faces (data not shown).

Following data pre-processing, one child participant was removed due to failure to acquire a complete 3D head digitisation (used for coregistration of the sensor locations to brain anatomy – see methods). We removed 13±9 (mean ± standard deviation) trials in children and 7±4 trials in adults due to excessive interference. Trials were then matched across age groups by randomly selecting and removing additional trials in adults and older children, this resulted in each age group having an average of 43 trials. On average we had 159±11 (mean ± standard deviation) channels of data at UoN, and 78±3 channels at SK. All data were processed using spatial filtering to derive images showing the spatial signature of task induced change in neural oscillations, and time-frequency representations of neurophysiological activity at locations of interest in visual cortex.

### Gamma oscillations change with age

Figures 2a-f, show the spatial and spectro-temporal signatures of gamma activity for all participants. Data were separated into six age groups and, for all groups, an image showing the spatial distribution of gamma modulation is shown (as a red overlay on the standard brain, averaged across subjects). Time-frequency-spectra (TFS) extracted from the location of peak gamma modulation are also shown. In the TFS, yellow indicates a task-induced increase in oscillatory amplitude relative to baseline, whereas blue indicates a decrease (baseline was measured in the -0.8 – -0.1 s window prior to stimulation onset).

**Figure 2.**
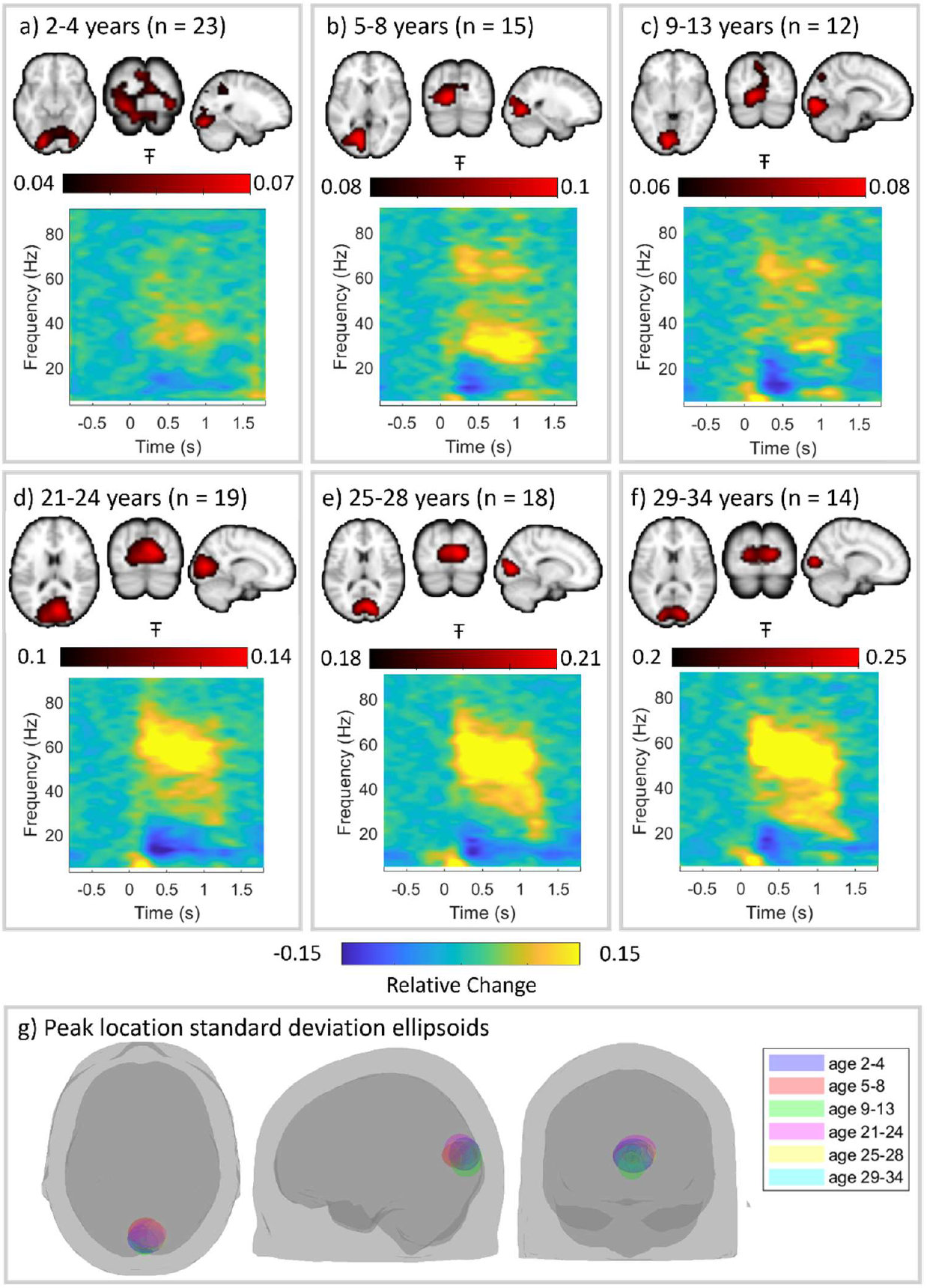
Age-group-specific time-frequency spectrograms show neurodevelopment of gamma oscillations. Participant averaged pseudo-T statical images of gamma modulation are shown in red (4mm resolution) overlaid on the standard brain. The time frequency spectrograms show group averaged oscillatory dynamics from the location of largest gamma modulation in visual cortex. a) 2–4-year-olds (n=23), b) 5–8-year-olds (n=15), c) 9–13-year-olds (n=12), d) 21–24-year-olds (n=19), e) 25–28-year-olds (n=18) and f) 29–34-year-olds (n=14) (ages are inclusive). Note the evolution of spectral signature with age. g) Ellipsoids describing the mean and standard deviation of the coordinates of the largest gamma modulation for all age groups. We saw no significant difference in the location of the visual gamma response with age in any axis (p=0.36, p=0.92 and p=0.52 for x,y and z axes, measured using Pearson correlation to test for a systematic shift in spatial localisation due to age).

We saw no significant difference in the location of the visual gamma response with age (see Figure 2g) in any axis. We did however see a changing spectro-temporal picture with age. In younger subjects we saw a task induced broadband gamma increase. As children age, the broadband response remains, and we also observed bimodal gamma activity, most prominent at around 35 Hz and 70 Hz. This further evolved to a broad band response with additional high amplitude narrow band activity at around 60 Hz in adults. This is consistent with previous literature (Bharmauria et al., 2016; Murty et al., 2018; Ray and Maunsell, 2011).

Figure 3 formalises the data in Figure 2 by demonstrating statistical significance of the observed spectral changes. The central graph shows stimulus induced relative change in oscillatory amplitude, for the 6 age groups, plotted against frequency. This was calculated by contrasting the 0.3 – 1 s window (during stimulation) to the -0.8 – -0.1 s (rest) window (Campbell et al., 2014). The inset plots show relative change in oscillatory amplitude, for individual participants, for frequency bands 11-15 Hz, 29-33 Hz and 51-55 Hz. Here, each data point represents a single individual in the study and data are plotted against age. Pearson correlation showed a significant (R = 0.57, *p* = 6.6 × 10^−10^) increase in spectral amplitude with age in the 51-55 Hz (gamma) range. There was no significant effect, however, at 11-15 Hz (alpha frequency range) or 29-33 Hz (low gamma) (R = −0.16, *p* = 0.1129 and R = −0.1, p = 0.3090, respectively). This is consistent with a stimulus induced broadband gamma increase at all ages, and emergent narrowband effects in adults.

**Figure 3.**
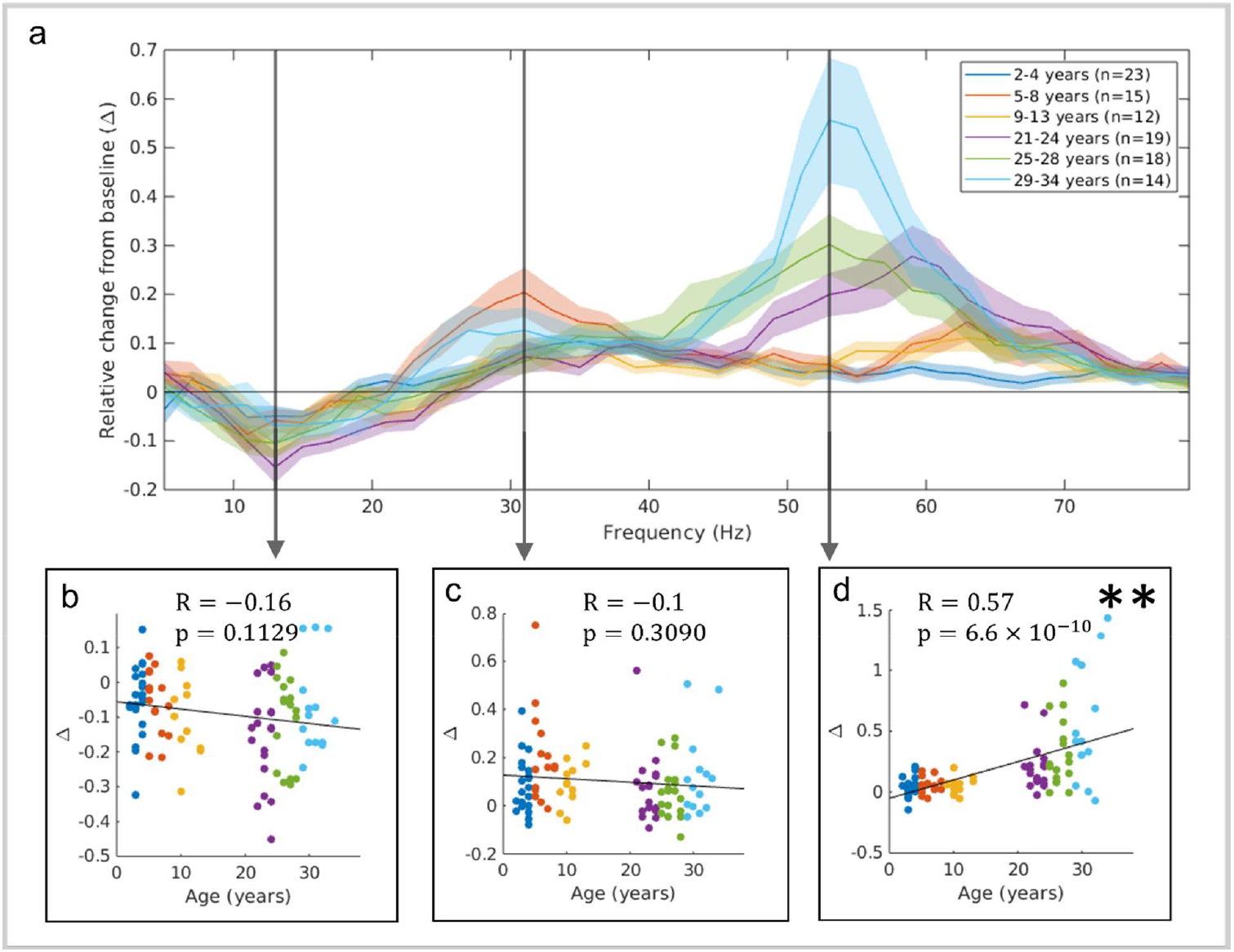
Gamma amplitude changes with age. The stimulus induced relative change in oscillatory amplitude from baseline is plotted against frequency for the 6 age groups (a). The relative change was measured in the 0.3 – 1 s window post-stimulus compared to the -0.8 s to -0.1 s baseline period. Lines show the group mean with shading representing standard error. The inset scatter plots (b, c and d) show relative change for all individuals in the study plotted against age (colour indicating age group), with straight lines fitted to the data. Specifically, we show data in the frequency ranges 11-15 Hz (b) (R = −0.16, p = 0.1129); 29-33 Hz (c) (R=-0.1, p = 0.3090) and 51-55 Hz (d) (R = 0.57, p = 6.6 × 10^−1^). The star (*) indicates uncorrected significance (p < 0.05) and (**) indicates significance following Bonferroni correction with a threshold of p < 0.0011 to account for 44 comparisons across different frequency bands.

### DCM suggests E-I balance drives spectral changes

A local spectral DCM, optimised for V1 (Shaw et al. 2017), was used to determine how inhibitory and excitatory activity drives the observed changes in gamma oscillations between children and adults. Briefly, the individual subject difference spectra (between task and rest) were extracted. The absolute values were derived and fitted to a model which optimises a set of parameters describing how contributions from different cellular assemblies, in different cortical layers, contribute to the observable signal. This model, which is summarised by Figure 4a, has been verified in recent literature using adult MEG recordings and pharmacological intervention (Shaw et al., 2020, 2017). Figure 4b shows the average (absolute) difference spectrum (between stimulation and rest) for all participants, highlighting the gamma change. Similar spectra (for individuals) were used to fit the DCM. A linear regression model, covarying for sex (Fung et al., 2021), was used to investigate the relationship between age and the model parameters. We also investigated the ratio between parameters in the superficial layer (G11/G12) to probe the hypothesized E-I balance specifically related to visual gamma (Shaw et al., 2017). Figure 4c shows the results, with significant age relations in parameters G5 (describing the excitatory output from spiny stellate cells to inhibitory interneurons) and the ratio between G11 and G12 (which represent the inhibitory and excitatory connections between superficial pyramidal neurons and inhibitory inter-neurons) following correction for multiple comparisons. Figure 4d shows scatter plots of model parameters (G5, G11/G12, and G11 and G12 individually) with age; notice that inhibition tends to increase, and excitation decrease.

**Figure 4.**
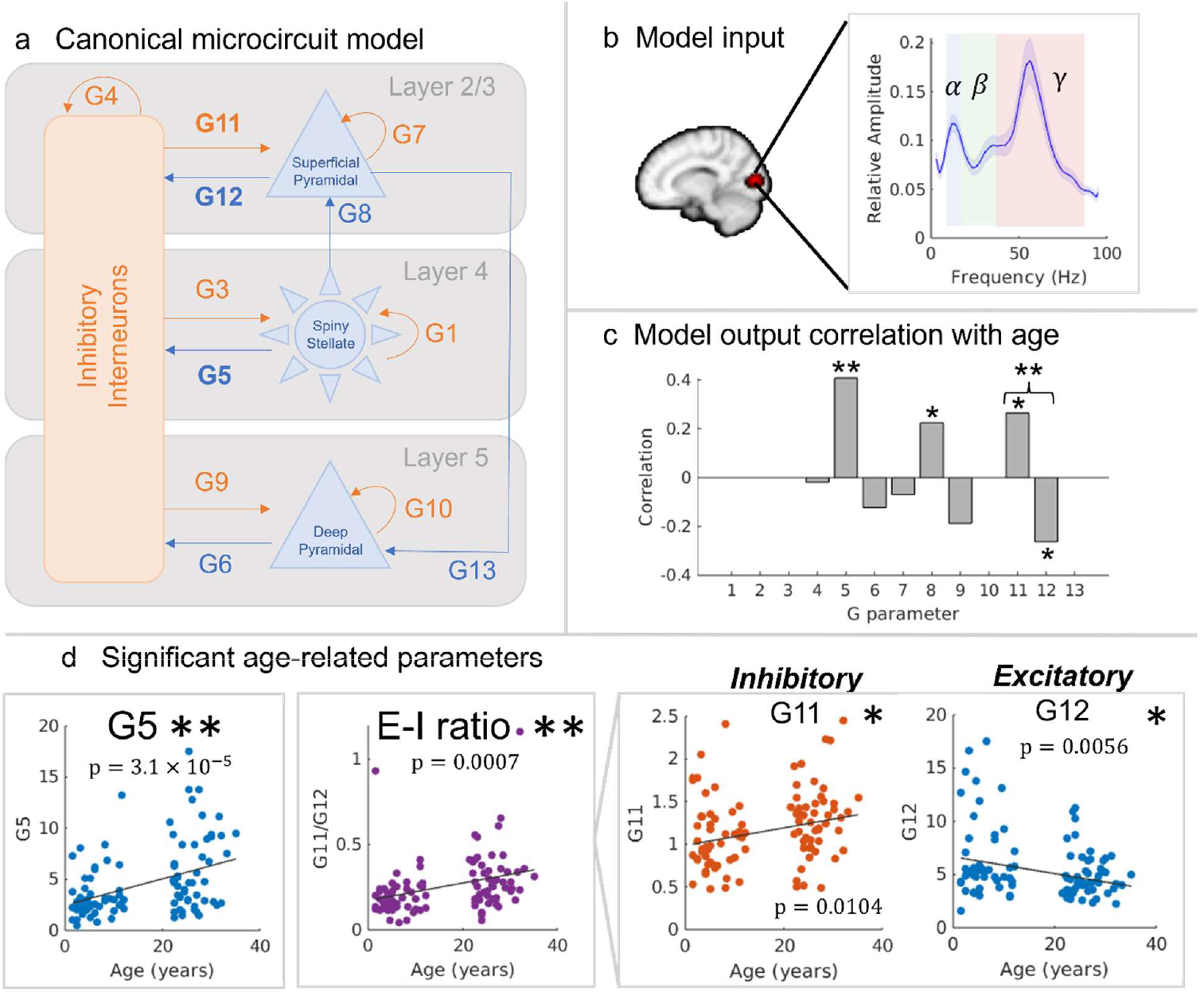
DCM suggests E-I balance underlies age related spectral differences. a) The canonical microcircuit model describes the relative contribution of cells within the cellular column. The model takes spectral input from data in visual cortex and fits a set of parameters (G1 – G12) which describe the relative contribution of the different neuronal assemblies to the measured signal. Excitatory signals are indicated by blue and inhibitory in orange. b) Average (across all subjects) absolute difference spectrum between active and control windows, with canonical frequency bands highlighted (alpha in blue, beta in green and gamma in red). c) Correlation of the model derived ‘G parameters’ with age. Significant age-relations were observed in G5 and the ratio of parameters G11 and G12. d) Scatter plots for G5 (excitatory); the E-I ratio of G11 and G12, and G11 (inhibitory) and G12 (excitatory) individually. The star (*) indicates uncorrected significance (p < 0.05) and (**) indicates significance following Bonferroni correction with a threshold of p < 0.0056 to account for 9 comparisons across parameters.

### Alpha suppression shifts in frequency with age

Finally, for completeness, we assessed how age affects stimulus induced change of alpha oscillations. Figure 5a shows the spatial signature of alpha suppression (in blue, overlaid on the standard brain) alongside the TFS data from the locations of largest task induced alpha modulation, across the age groups. Note that these regions differ from those of maximum gamma change, and consequently the gamma change is less prominent. Note that alpha modulation is clear in all groups.

**Figure 5.**
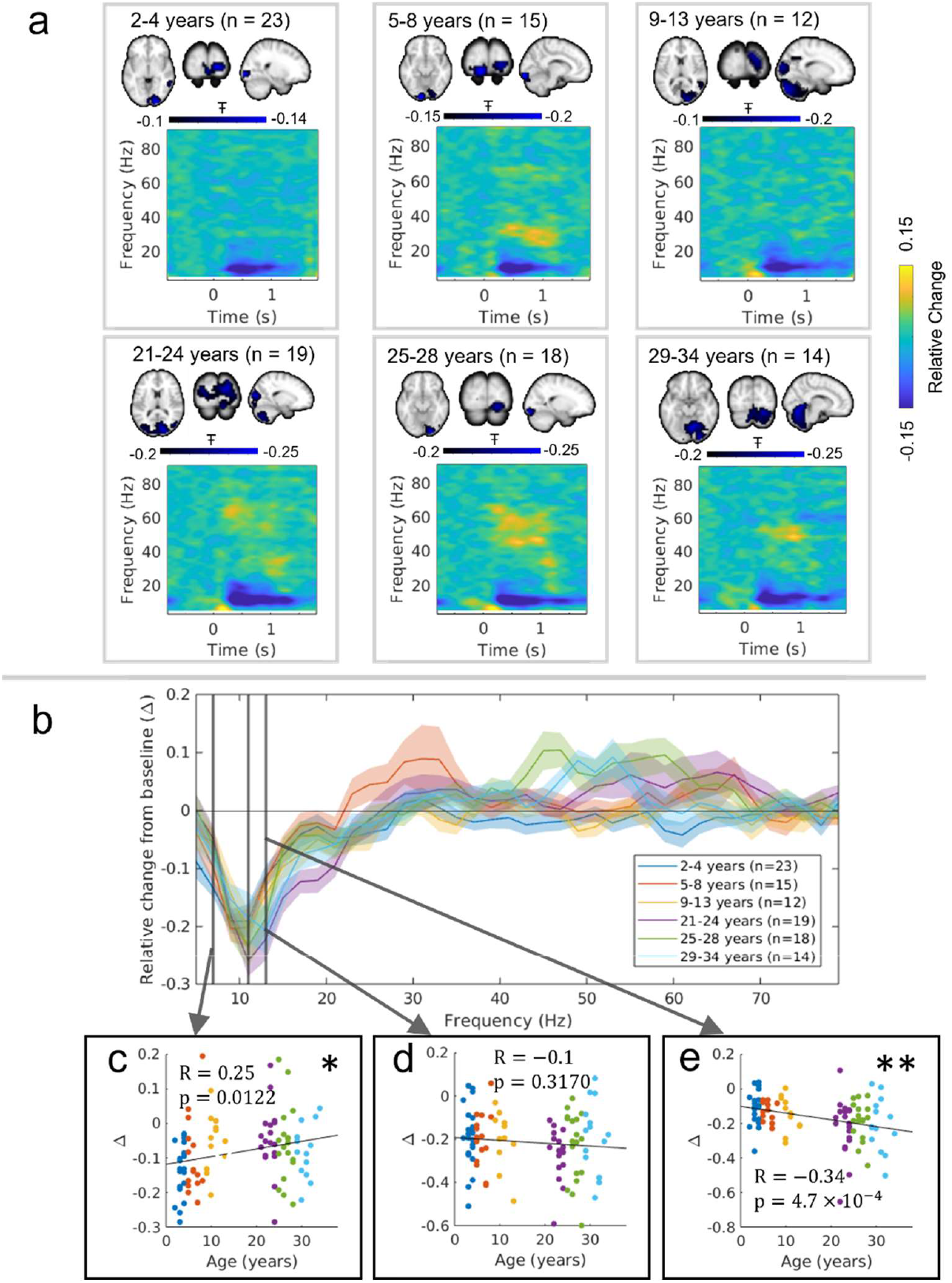
Alpha suppression remains across ages. a) Pseudo-T statistical maps and time-frequency spectrograms from the locations of peak of alpha suppression. Data are divided by age group. b) Relative change in oscillatory amplitude as a function of frequency. The inset scatter plots show how stimulus induced amplitude change differs for individuals in the c) 5-9 Hz range (R = 0.25, p = 0.0122), d) 9-13 Hz range (R = −0.1, p = 0.3170 and e) 11-15 Hz range (R = −0.34 p = 4.7 × 10^−^) bands (colour indicating age group). Adults show a significantly larger alpha suppression in the 11-15 Hz range. This is consistent with alpha modulation being lower frequency in younger participants. The star (*) indicates uncorrected significance (p < 0.05) and (**) indicates significance following Bonferroni correction with a threshold of p < 0.0011 to account for comparison across 44 frequency bands.

In Figure 5B, the spectrum shows relative change in oscillatory amplitude from baseline as a function of frequency (including a zoomed in area over the alpha band). The inset scatter plots show relative change, for individual participants, for the frequency bands 5-9 Hz, 9-13 Hz and 11-15 Hz. We found no change in alpha modulation for the 9-13 Hz canonical alpha band. However, we saw increased (more negative) 5-9 Hz modulation in younger participants (though this was non-significant following correction for multiple comparisons) and increased 11-15 Hz modulation for older participants. This is in broad agreement with the widespread finding that the alpha rhythm’s peak frequency tends to increase with age (Miskovic et al., 2015).

## Discussion

E-I balance (or imbalance) underpins healthy and atypical brain function and its characterisation could provide useful insights into neurodevelopmental disorders (Sohal and Rubenstein, 2019). While in-vitro and animal studies form the basis of such models, the ability to non-invasively characterise E-I balance using imaging offers a means to bridge the gap between experimental animal and in-vivo human physiology. Gamma oscillations provide a window on E-I balance, yet the formation and developmental trajectory of gamma oscillations in humans, through the early years of life, remains poorly understood. This study is the first to capitalize on the potential of OPM-MEG for the investigation of gamma oscillations from toddlerhood to adulthood, and the first to apply a DCM to OPM data to explore the neurochemical underpinnings of gamma signals.

Using a well-established visual paradigm, we showed that age has a significant impact on the spectro-temporal neurophysiological response from visual cortex. In the broadband gamma frequency range (30-80 Hz), low-amplitude oscillations are present, even in early life and appear to remain to adulthood. However, in later childhood we see a multi-spectral response, followed by higher-amplitude band limited oscillations (at ∼60 Hz) which emerge in adulthood. This latter finding (in adults) is in strong agreement with previous studies (Hoogenboom et al., 2006; Muthukumaraswamy et al., 2010). Statistical analyses showed a significant increase in oscillatory amplitude with age in the 51 – 55 Hz window. Despite these significant spectral changes, we saw no measurable shift in the spatial origin of gamma oscillations with age, with the maximum signal consistently localised to primary visual cortex. Our results also highlight that visual gamma, even in adults, has high inter-individual differences and this agrees with other studies employing similar paradigms (e.g. (Muthukumaraswamy et al., 2010)). We examined suppression of alpha amplitude during visual stimulation, which was relatively stable across age groups. In the 9 – 13 Hz band, alpha suppression showed no significant relationship with age; this provides a key validation of data quality across our dataset (i.e. if data were of poorer quality in younger participants, we would likely see a drop in alpha suppression in those individuals, which is not the case). We did however see a trend towards increased 5-9 Hz modulation in younger participants, and significantly increased 11-15 Hz modulation in adults. This is in good agreement with other studies (Miskovic et al., 2015) which show a shift in alpha peak frequency with age (albeit typically in resting state data), with younger subjects tending to have a lower alpha frequency. This provides further verification of our data quality.

Our DCM shows how age-related changes in gamma oscillations are driven by a neural circuit that matures with age. Results suggest that several parameters demonstrate an age dependency. Specifically, excitatory signals from spiny stellate cells to inhibitory interneurons (parameter G5), and the relative inhibitory vs. excitatory signalling from superficial pyramidal neurons to inhibitory interneurons (the ratio of parameters G11 and G12) both significantly increased in adults compared to children. Previous work has demonstrated that G5 relates to beta and gamma amplitudes (Shaw et al. 2017) and so this is in strong agreement with our spectral results, where we showed increased gamma amplitude in older participants. An increase in the ratio between G11 and G12 supports our initial hypothesis that maturation would see a change in E-I balance (Larsen et al., 2022), such that inhibition in the superficial layer of the visual cortex increases, while excitation decreases, as children grow up. This is likely due to an increase in gamma aminobutyric acid (GABA) (Jansen et al., 2010) and a relative decrease in glutamate (Hädel et al., 2013). We are the first to implicate these age-related changes via assessment of visual gamma oscillations.

This study provides an important foundational step in the measurement of E-I balance via gamma oscillations in neurodevelopment. However, there are limitations which should be addressed. Firstly, OPM-MEG systems remain new technology; OPMs have a higher noise floor than conventional MEG sensors, and the number of measurement channels is lower (again compared to conventional MEG instrumentation). However, we did use helmets which are lightweight, allow subject movement, and come in multiple sizes enabling adaptation for age. This alleviates confounds of SNR change with age and movement – which (anecdotally) was large in children. We believe this study would not have been possible using either conventional MEG (due to confounds of head size and movement) or EEG (due to gamma oscillations being obfuscated by muscle artifacts). Importantly, OPM systems are still under development, and it is highly likely that sensor density (Hill et al., 2024) and noise floor will improve with time, meaning OPM-MEG will likely become the technique of choice for high-fidelity characterisation of brain function in neurodevelopment in the future. Secondly, to increase participant numbers, data were collected from two sites, potentially introducing a confounding effect of scanner configuration. To mitigate this, we matched recording conditions as far as possible, and a cross-site comparison within our adult groups (Figure S1) showed no significant differences between sites. Further, at both sites we studied children and adults, meaning any measurable age-related differences are unlikely to be driven by site. We, therefore, think it unlikely that our results could be affected by the cross-site nature of recordings; indeed, the fact that we were able to demonstrate cross-site reliability is extremely positive to accelerate the (already rapid) uptake of OPMs and to support the collection of new large, across-site datasets. A final limitation is that we have a non-uniform range of participant age; whilst this was enough to demonstrate significant age-related changes, the addition of adolescents and older adults to this study would likely enable elucidation non-linear trajectories. Future work should aim to fill these gaps.

An imbalance in excitatory and inhibitory neurotransmission underlies current theories for the pathophysiological underpinnings of neurodevelopmental and psychiatric disorders. However, the study of these signals has been limited by technology, restricting most studies to adults, animal models and the lab benchtop. OPM-MEG lifts these constraints, allowing us to measure signals relating to E-I balance directly, and from early life. We have demonstrated this important milestone and our results – which show significant changes in gamma oscillations and E-I balance with age - offer insight into early cortical maturation and provide a typically developing standard, from which clinical applications can be explored.

## Online Methods

### Data Acquisition

The UoN OPM array comprised 64 triaxial OPMs (3^rd^ generation QZFM; QuSpin, Colorado, USA) enabling up to 192 channels of magnetic field measurement. The SK system comprised 40 dual-axis OPMs (3^rd^ generation QZFM; QuSpin), enabling up to 80 channels of magnetic field measurement.

In both systems, sensors were combined to form an array and integrated with other hardware (e.g. for magnetic field control) and software (e.g. for stimulus delivery and data acquisition) to form two complete MEG systems (Cerca Magnetics Ltd, Nottingham UK). Specifically, sensors were mounted in rigid 3D-printed helmets (five sizes were available). Participants wore a thin aerogel cap or had insulating padding under the helmet for thermal insulation. Participants were seated in a patient support at the centre of a magnetically shielded room (MSR). The UoN system was housed in an OPM-optimised MSR which comprises 4 layers of mu-metal, one layer of copper, and is equipped with degaussing coils. The SK system was housed in a repurposed MSR from a cryogenic-MEG system which comprised two layers of mu-metal and one layer of aluminium (Vacuumschmelze, Hanau, Germany). In both systems, bi-planar coils (Cerca Magnetics Limited) surrounded the participants to provide active magnetic field control (Holmes et al., 2018). In the UoN instrument, coil currents were applied to cancel out the residual (temporally static) magnetic field (Rea et al., 2022; Rhodes et al., 2023; Rier et al., 2024). At SK (where time-varying field shifts were larger) a reference array provided dynamic measurement of the environmental magnetic field and feedback to the bi-planar coils enabled real-time compensation of both static and dynamic magnetic field changes (Holmes et al., 2019). Equivalent data from these two systems have been demonstrated previously (Hill et al., 2022). In both systems, participants were free to move throughout data acquisition (but were not encouraged to do so). Data were collected at a sampling rate of 1200 Hz, from all sensors, using a National Instruments (NI, Texas, US) data acquisition system interfaced with LabView (NI).

For coregistration of sensor geometry to brain anatomy, two 3D digitisations of the participant’s head (with and without the OPM helmet) were acquired using a structured light camera (Einscan H, SHINING 3D, Hangzhou, China). These digitisations, coupled with accurate knowledge of the helmet structure from its computer aided design (CAD) allowed knowledge of the sensor locations/orientation relative to the head. They also enabled generation of a ‘pseudo-MRI’ which provided an approximation of the underlying brain anatomy (for more details see (Rier et al., 2024)). Briefly, age-matched template MRIs (Richards et al., 2016) were warped to the individual participant’s 3D head digitisation using FSL FLIRT (Jenkinson et al., 2002). For some of the youngest participants, head digitisation failed and so the age-matched templates were used as the pseudo-MRI without warping.

### Participants and Paradigm

The study was approved by the local research ethics board committee at both sites. All adult participants provided written informed consent. A legal guardian for all participants under 18 years provided the written informed consent and the child gave verbal assent. 27 children and 26 adults took part in the study at UoN; 24 children and 26 adults were scanned at SK. Children were always accompanied by a parent and at least one experimenter inside the MSR. Adult data were sex- and age-matched across the two sites to enable a cross-site comparison.

Visual stimulation comprised an inwardly moving drifting circular grating. The grating was displayed centrally and subtended a visual angle of 7.6°. A single trial comprised 1000 ms of stimulation followed by a jittered rest period of 1250 ± 200 ms. 60 trials in total were shown and these ‘circles’ trials were interspersed with images of faces (data not included). Precise timing of the onset and offset of stimulation was sent from the stimulus PC to the OPM-MEG system via a parallel port.

### Data Analysis

Data processing was identical at both sites. Bad channels (those that either had high noise or zero signal) were identified by manual inspection of the channel power spectra and removed. Data were notch filtered at the powerline frequency (50 Hz for UoN and 60 Hz for SK) and 2 harmonics. A 1 – 150 Hz band pass filter was applied, following which, data were epoched to 3 s trials encompassing 1 s prior to the onset of the “circle” and 2 s after. Bad trials were identified as those with trial variance greater than 3 standard deviations from the mean and were removed. Visual inspection was carried out and any further trials with noticeable artefacts were removed. ICA was used to remove eye blink and cardiac artefacts (implemented in FieldTrip (Oostenveld et al., 2011)) and homogeneous field correction (HFC) was applied to reduce interference that manifests as a spatially homogeneous field (Tierney et al., 2021).

We used an LCMV beamformer to project magnetic fields recorded at the sensors into estimates of current dipole strength in the brain (Van Veen et al., 1997). The forward model was constructed using a single-shell model (Nolte, 2003), fitted to the pseudo-MRI and implemented in FieldTrip (Oostenveld et al., 2011). Voxels were placed on an isotropic 4-mm grid covering the whole brain, and an additional 1-mm isotropic grid covering the visual cortex (identified by dilating a mask of the left and right cuneus from the AAL atlas (Hillebrand et al., 2016; Tzourio-Mazoyer et al., 2002) with a 5 mm spherical structuring element). Covariance matrices were generated using 1-150 Hz broadband data spanning all circles trials (excluding bad trials), regularized using the Tikhonov method with a regularization parameter of 5% of the maximum eigenvalue of the unregularized matrix (Brookes et al., 2008). This matrix was used to compute the beamformer weighting parameters used for all subsequent calculations.

Pseudo-T statistical images were constructed by contrasting either alpha or gamma power during stimulation and rest. Specifically, we derived four additional covariance matrices (*C*_*ON_alpha*_, *C*_*OFF_alpha*_, *C*_*ON_gamma*_ and *C*_*OFF_gamma*_). For the gamma matrices, we used 30 – 80 Hz filtered data and for alpha band we used 6 – 14 Hz filtered data. The ON window was 0.3 – 1 s and the OFF window was -0.8 – -0.1 s (timings relative to the onset of the circle.

TFSs showing neurophysiological activity at the locations of maximum gamma/alpha modulation (identified using the 1-mm resolution images) were derived. TFS data in the 1 – 100 Hz frequency range were generated by first sequentially filtering broadband beamformer projected data into 45 overlapping frequency bands (2 Hz separation, 4 Hz bandwidth). For each band, the Hilbert transform was computed to give the analytic signal; the absolute value was computed to derive a measure of instantaneous oscillatory amplitude, and these ‘Hilbert envelopes’ were averaged across trials and concatenated in the frequency dimension. For each band, a mean baseline amplitude was taken (in the -0.8 s to -0.1 s) window and subtracted. Data were then normalised by the baseline values to give a measure of relative change in amplitude. These data were collapsed in time to give spectral relative change (i.e. Figures 3 and 5). In all cases, we investigated the statistical relations between age and amplitude modulation using Pearson correlation.

#### DCM

Neurophysiologically informed modelling was performed using dynamic causal modelling (DCM) for steady-state responses implemented in SPM8 (Moran et al., 2009; Shaw et al., 2017). The canonical microcircuit structure (CMC, shown in Figure 4a) describes a simplified model that strikes a balance between biological reality and complexity that can be modelled. The model estimates membrane potentials and postsynaptic currents of cell populations through differential equations. We differ here from the analysis described in Shaw et al. 2017 by using relative spectra rather than pre-whitening by removal of the 1/f profile, as this proved advantageous for OPM data, where absolute spectra are more prone to noise (see also Discussion). Relative spectra from the beamformer estimated time series at the peak gamma modulation were calculated by taking the power spectral density (PSD) of data during the stimulus (0.3 – 1 s) minus the PSD of data during the rest (−0.8 to -0.1 s) windows. Spectra are normalised such that the area under the global average equals 1, but relative peak height is preserved. Model priors are determined from the global average and parameters that have little or no effect (G1, G3, G10 and G13) are held constant as in prior work (Shaw et al., 2017). Finally, we separately model the alpha peak frequency using a single Gaussian (constrained to 8 to 13 Hz) and remove this, as the model is capable of generating clear beta and gamma peaks but alpha is thought to be generated over more extensive circuity (Bastos et al., 2014). These processes allow the DCM to estimate the G parameters that result in the measured beta and gamma responses.

## Data and code availability

Data from UoN will be made available on Zenodo. Data from SickKids will be available through Ontario Brain Institute. OPM analysis code will be made available on GitHub (https://github.com/nsrhodes/gamma_opm_2024). Dynamic causal modelling was performed using a variant of DCM-SSR in SPM8 and code will be made available upon request.

## Author contributions

N.R.-research design, data collection, data analysis, data interpretation, writing paper, L.R.-data collection, data interpretation, writing paper – reviewing and editing, K.S. – data analysis, data interpretation, writing paper – reviewing and editing, J.S. – data collection, data interpretation, writing paper – reviewing and editing, M.M.V. – data analysis, data interpretation, writing paper-reviewing and editing, N.H. – system fabrication, data collection, data interpretation, writing paper-reviewing and editing, E.B. - system fabrication, writing paper-reviewing and editing, R.M.H. – system fabrication, writing paper-reviewing and editing, M.J.T. – supervision, funding acquisition, research design, data interpretation, writing paper, M.J.B. – supervision, funding acquisition, research design, data interpretation, writing paper

## Acknowledgements

We would like to thank all our participants and the families of our young participants for their involvement in this study.

