## Supplementary information for "Measuring the neurodevelopmental trajectory of excitatory-inhibitory balance via visual gamma oscillations"

**Supplementary Information Text**

**Additional Methods**

**Site comparison:**

A table describing the differences between the two systems is shown in Table. S1.

| | **UoN** | **SickKids** |
|---|---|---|
| **No. sensors** | 64 triaxial | 40 dual-axis |
| **No. channels** | 192 | 80 |
| **Helmet** | Rigid 3D printed helmet in one of three sizes (adult S, adult L, 4-year-old) | Rigid 3D printed helmet in one of two sizes (adult L, 4-year-old) |
| **Magnetic environment** | Quiet campus location outside of the city centre | Downtown location above a busy car park near construction sites, main roads and underground transport lines |
| **Passive shielding** | OPM-optimised shielded room with four layers of mu-metal and one layer of copper | Repurposed room with two layers of mu-metal and one layer of aluminium |
| **Degaussing coils** | Yes | No |
| **Magnetic environment with passive shielding** | ~ 2 nT static magnetic field with drifts of $\pm$ 1 nT across 10 minutes (Rea et al., 2021) | ~ 30 nT static magnetic field with drifts of $\pm$ 20 nT across 10 minutes (Hill et al., 2022) |
| **Active shielding (static)** | Via average coil currents through biplanar coils | Via reference array through biplanar coils |
| **Active shielding (dynamic)** | Not required (due to minimal dynamic drift) | Via reference array through biplanar coils |

***Table S1. Site comparison.*** *A table describing the features of the OPM-MEG systems at each site.*

There are three main differences between the two systems. The first of which is the number of channels of MEG data. The UoN system consisted of 64 triaxial OPM sensors (QuSpin Inc. Triaxial Gen-3), providing up to 192 channels of data. The SickKids system used 40 dual axis OPM sensors (QuSpin Inc. Dual-axis Gen-3), providing up to 80 channels of data. It is important to note that the radial axis (available with both Dual- and tri-axial OPMs) provides the largest MEG signal (Iivanainen et al., 2017; Sarvas, 1987) and as such, the increase in sensitivity to the MEG signal is not directly related to channel count with triaxial sensitivity. A second difference is the magnetic environment of each site, with a contrast in location between a city-centre lab and a green campus-based lab, as described by Hill et al. (Hill et al., 2022). Finally, the magnetic shielding between systems is different, with UoN benefitting from OPM-optimized shielding whereas the SickKids system uses a re-purposed room from a cryogenic MEG system. To summarize, despite differences between the two OPM-MEG sites and systems, we found no significant differences in visual gamma response between our age- and sex-matched groups of healthy adults (see Figure S1). This builds upon the result from a previous study with an individual participant (Hill et al., 2022), extending the comparison to consider group analyses.

**The "Faces Circles" paradigm:** A diagram of the paradigm is shown in Figure 1. b) shows an example of the circular grating image, with arrows depicting the direction of travel of the circular grating. Images of emotional faces are also shown, with trials interspersed between circles trials but were not analyzed in this study. Additional cartoon character catch trials were used to maintain attention from young participants. Each trial started with a white fixation cross located centrally on a black screen, presented for an interstimulus interval (ISI) randomly jittered between 1050 – 1250 ms. Following this, either an emotional face image was presented for 500 ms, or an inward drifting circular grating (oscillating at 1.2 $°s^{-1}$) was presented for 1 s. This was repeated up to a total of 60 circles trials. There were between 120 – 160 faces trials due to UoN running a version of the paradigm that included an additional emotion (but these data were not analyzed in this work). The total experiment lasted between 5-8 minutes (due to the varying number of face trials).

**Participant demographics:** Table S1 provides a detailed description of the demographics from each group of participants, separated into UoN and SickKids adults and children. One child was excluded from analyses due to the inability to acquire a 3D digitization for coregistration. In addition, one adult participant scan from SickKids was excluded from the main text analyses due to repeat measures to result in 101 unique participants. The repeat scan was included in the cross-site comparison in Figure S1.

| | No. scans | Age (years, mean +/- standard deviation) | Age range (years) | No. males | No. females |
|---|---|---|---|---|---|
| **UoN Children** | 27 | $7.67 \pm 3.09$ | 2 – 13 | 10 | 17 |
| **UoN Adults** | 26 | $26.35 \pm 3.29$ | 21 – 34 | 12 | 14 |
| **SickKids Children** | 24 | $3.79 \pm 0.76$ | 3 – 5 | 10 | 14 |
| **SickKids Adults** | 26 | $26.27 \pm 3.26$ | 21 – 34 | 12 | 14 |
| Total | 103 | $16.17 \pm 10.71$ | 2 – 34 | 44 | 59 |

***Table S2. Participant demographics.*** *A table describing the demographics of the participants scanned in the study across the two sites and age group cohorts.*

### Additional Results

**Cross-site comparison:** Figure S1 shows a comparison of gamma measurements acquired at the two sites. Data are shown for two age and sex-matched adult groups. Figure S1a shows the mean and standard deviation of the locations of the peak change in gamma modulation; these locations have been derived for each participant, projected to an average brain, and are shown as ellipsoids (i.e., the axes of the ellipsoid represent the standard deviation of the peak location coordinates in x, y and z). We show results for three cases; UoN all channels; SK all (80) channels and UoN 80 channels; the latter is included to ensure equivalence in terms of channel count across sites. We found no significant differences between peak locations, suggesting that the two systems were equivalent in terms of delineation of the spatial signature of gamma modulation.

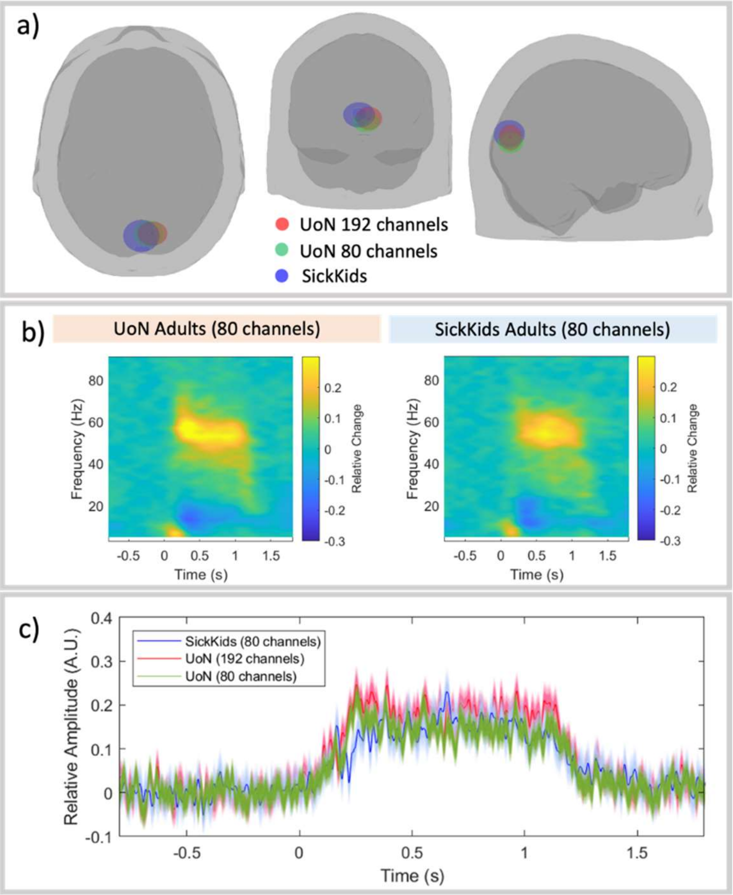


***Figure S1. Cross-site comparison in a group of adults.*** *a) The mean and standard deviation of peak locations for gamma modulation. The centre of the ellipsoids represents the mean location of maxima; the ellipsoid axes represent standard deviation in the x, y and z coordinates. The UoN analyses performed for both the full channel count and a reduced channel count (to 80) to match the SK system. b) The time-frequency spectrograms extracted from virtual electrodes at the peak of gamma modulation. c) Gamma band envelopes across groups at SK and UoN, again with the UoN analyses performed for both the full channel count and reduced channel count.*

Figure S1b shows the group averaged time frequency spectra (TFS) for participants scanned at UoN (left) and SickKids (right). In both cases, the TFS is derived from a virtual electrode at the location of maximum gamma modulation; blue represents a decrease in oscillatory amplitude relative to baseline, yellow an increase. Time zero represents stimulus onset, and here, the UoN system has been reduced to 80 channels for comparison with SK. As expected, we see large increases in gamma activity with concomitant decreases in alpha and beta oscillations. Most importantly activity at both sites looks similar. Figure S1c shows the envelope of 30 – 80 Hz oscillatory amplitude for the UoN system with full channel count (up to 192 channels) in red, UoN reduced channel count in green and SickKids system in blue. A statistical analysis showed no significant differences between systems when measuring relative change in amplitude (measured between active ($0.3 \text{ s} \leq t \leq 1 \text{ s}$) and baseline ($-0.8 \leq t \leq -0.1 \text{ s}$) windows; $T = 1.01$, $p = 0.32$ for the full UoN channel count; $T = -0.26$, $p = 0.80$ for the reduced channel count). These data suggest no measurable difference between sites when assessing the temporal modulation of gamma activity.

**Complete statistics from cellular modelling:** Table S3 provides the Pearson correlation and Bonferroni-corrected p-values for the relation between the spectral parameters of the data and the corresponding G parameters from the model.

| | G4 | G5 | G6 | G7 | G8 | G9 | G11 | G12 |
|---|---|---|---|---|---|---|---|---|
| **Beta frequency** | 0.08 [0.4059] | -0.08 [0.4329] | 0.41 [2.6e-5] | 0.05 [0.6817] | 0.14 [0.1649] | 0.00 [0.9905] | 0.16 [0.1168] | 0.15 [0.1250] |
| **Beta amplitude** | -0.03 [0.7829] | 0.40 [3.7e-5] | 0.42 [1.3e-5] | 0.08 [0.4067] | 0.44 [5.8e-6] | 0.47 [8.6e-7] | 0.29 [0.0031] | 0.19 [0.0614] |
| **Gamma frequency** | 0.03 [0.7935] | -0.26 [0.0091] | -0.04 [0.6852] | 0.19 [0.0625] | -0.27 [0.0072] | 0.05 [0.6109] | -0.36 [2.6e-4] | -0.02 [0.8735] |
| **Gamma amplitude** | 0.02 [0.8426] | 0.66 [4.8e-14] | 0.17 [0.0876] | -0.05 [0.6506] | 0.58 [1.9e-10] | -0.13 [0.1824] | 0.49 [2.0e -7] | -0.16 [0.1082] |

***Table S3. G parameters and MEG features.*** *A table outlining the Pearson correlation [with p-value] between spectral features of the MEG signal and the model parameters. Significant relations are highlighted in red (positive) and blue (negative). The threshold for significance is corrected for multiple comparisons using Bonferroni correction and set as 0.0016 (32 comparisons).*

Table S4 provides the summary of the statistics from the dynamic causal model describing the relation between G parameters and age while covarying for sex.

| | G4 | G5 | G6 | G7 | G8 | G9 | G11 | G12 | G11:G12 |
|---|---|---|---|---|---|---|---|---|---|
| **R** | -0.001 | 0.129 | -0.018 | -0.012 | 0.033 | -0.031 | 0.010 | -0.079 | 0.005 |
| **Standard error** | 0.006 | 0.030 | 0.015 | 0.020 | 0.014 | 0.017 | 0.004 | 0.028 | 0.002 |
| **T statistic** | -0.161 | 4.372 | -1.239 | -0.625 | 2.383 | -1.786 | 2.613 | -2.833 | 3.500 |
| **p value** | 0.8723 | 3.1e-5 | 0.2181 | 0.5332 | 0.0191 | 0.0772 | 0.0104 | 0.0056 | 0.0007 |

***Table S4. G parameter correlation with age.*** *Statistical analysis of the relation of model outputs with age, when covarying for sex. The threshold for significance is corrected for multiple comparisons using Bonferroni correction and set as 0.0056 (9 comparisons).*
